# Cardiac adenylyl cyclase type 8 overexpression increases locomotor activity in mice by modulating EEG-gamma oscillations

**DOI:** 10.1101/2021.07.19.452502

**Authors:** Jacopo Agrimi, Danilo Menicucci, Jia-Hua Qu, Marco Laurino, Chelsea D Mackey, Laila Hasnain, Yelena S Tarasova, Kirill V Tarasov, Ross A McDevitt, Donald B Hoover, Angelo Gemignani, Nazareno Paolocci, Edward G Lakatta

## Abstract

The central nervous system modulates heart function on a beat-to-beat basis via increasingly understood mechanisms. Conversely, whether and how humoral/functional cardiac variations shape brain activity and adaptive behavior remains unclear. This study shows that mice overexpressing adenylyl cyclase type 8 in myocytes (TGAC8), characterized by persistently elevated heart rate/contractility, also display increased locomotion. This effect is sustained by enhanced gamma rhythms, as evidenced by simultaneous behavioral and EEG/ECG monitoring. These changes are specific because they are not paralleled by other modifications, such as heightened anxiety-like behavior. In unison, TGAC8 mice hippocampus exhibits upregulated GABA-A receptors, whose activation chiefly accounts for gamma activity generation. Moreover, the Granger causality analysis between ECG and EEG attests to the causal involvement of the autonomic component of the heartbeat in shaping EEG gamma oscillations in a bottom-up modality. Mechanistically, TGAC8 harbors elevated circulating dopamine/DOPA levels of cardiac origin and upregulated hippocampal D5 dopamine receptor levels. In synergy with the GABA-A receptor, D5 activation favors hippocampal inhibitory currents that drive EEG gamma oscillations. These studies, therefore, inform how heart-initiated functional and/or humoral modifications reverberate back to the brain to modulate specific primary adaptive responses, such as locomotion.

**Significance:** The brain is continuously aware of the functional status of many bodily organs, modulating, for instance, the heart’s activity beat-by-beat. Conversely, how cardiac activity modifications impact brain function and behavior is less understood. We disclose that augmenting myocyte adenyl cyclase 8 (AC8) activity in mice increases their locomotion. Elevated cardiac AC8 levels lead to higher circulating dopamine and DOPA, hormones crucially involved in movement control, and increased expression of the hippocampus’s GABA-A and D5 receptors; the activation of the latter modifies hippocampal gamma oscillations shaping locomotor activity. Thus, the brain interprets changes in myocardial AC8 activity as a “sustained exercise-like” situation and responds by activating areas commanding to increase locomotion.

## Introduction

Damasio’s somatic theory of emotions postulates that afferent somatic signals from the body’s peripheral districts are integrated into subcortical and cortical regions, shaping emotional and behavioral responses needed to restore the body’s homeostasis (1). At the same time, however, the current perception of stress has significantly departed from the old idea of stress biology as “relevant only under unusual and threatening conditions”(2), in favor of the new notion of “an ongoing, adaptive process of assessing the environment”. The individual’s ability to sense changes intrinsic and extrinsic to the body would enable her/him to anticipate and thus cope better with future challenges (2).

At close inspection, the bidirectional relationship between the heart and the brain falls entirely within this loop. Indeed, besides the autonomic nervous system’s descending control of cardiac function, information is also processed by the intrinsic cardiac nervous system (the so-called “little brain of the heart”) that communicates back to the brain via ascending fibers located in the spinal cord and vagus nerve (3). These afferent impulses reach relay stations such as the medulla, hypothalamus, thalamus, and, ultimately, the cerebral cortex, carrying sensory information (3). However, despite numerous observational clues supporting the possibility that changes in cardiac activity can anticipate future challenges, thus alter behavior (4–6), definitive mechanistic evidence validating this view and putting forth pathways potentially accounting for these *bottom-up* phenomena remains to be gained.

Mice harboring a cardiac-specific overexpression of adenylyl cyclase (AC) type 8 (TGAC8) display a persistently elevated heart rate (HR), reduced HR variability (HRV), and increased contractility owing to enduringly elevated cardiac-intrinsic cAMP-PKA-Ca^2+^ signaling (7, 8). Moreover, the heart of these transgenic mice escapes top-down autonomic surveillance by blocking beta-adrenergic signaling and catecholamine production to escape harmful additional sympathetic stress (7). Further, the endocrine portfolio of the TGAC8 heart showcases an accentuated dopamine anabolism mirrored by elevated circulating levels of dopamine and DOPA (7), and the latter can permeate the blood-brain barrier (9). With this tool and evidence in hand, in the present study, we sought to model Damasio’s theory of the somatic marker in an experimental setting, proving that bioelectrical, mechanical, and endocrine signaling, borne in the periphery, can shape the activity of specific brain areas and consequently, adaptive behaviors.

## Results

### Mice with cardiac overexpression of AC8 display increased locomotor activity

Whether chronically elevated heart rhythm and/or contractility impact fundamental brain functions is currently unclear. Hence, we first set out to determine whether persistently elevated cardiac chronotropy/inotropy emanating from AC8 overexpression would affect two main behavioral variables, i.e., locomotion and exploratory behavior (i.e, anxiety/like behavior): two major behaviors associated with heart rate modifications (10). When challenged with an open field (OF) test, TGAC8 mice traveled more distance and at a higher speed than their age-matched littermates (+43% and +38%, respectively, p<0.01) (Fig. 1 B, C). Next, we determined their exploratory activity and anxiety-like behavior using the elevated plus maze (EPM) test. First, the EPM test confirmed the ability of TGCA8 mice to cover a longer distance at a higher average speed (Fig.1H, I), and with a maximum speed (Fig.1J) sizably higher than in the controls. Second, this test revealed a considerable decrease in time and episodes of freezing behavior in the transgenic mice (−45%, *p<0.05) (Fig. 1K, L). However, TGAC8 and WT did not differ in the two main parameters evaluating anxiety-like behavior, i.e., the number and times of entry in the open arms (Fig. E, F), results confirmed by the light-dark box test (Sup. Mat.). Further to unveil a possible anxious behavior, we performed the fear conditioning test, which, in contrast to the EPM, cannot be biased by concomitant alterations in locomotor activity (11). Thus, there was no sizable difference between the WT and TGAC8 on the basis of these two main trial parameters, time freezing contextual and cued (Fig. M, N). Hence, these findings indicate that TGAC8 mice - harboring persistently elevated chronotropic/inotropic activity at the heart level – display a hyperkinetic behavior without corollary anxiety.

**Figure 1.**
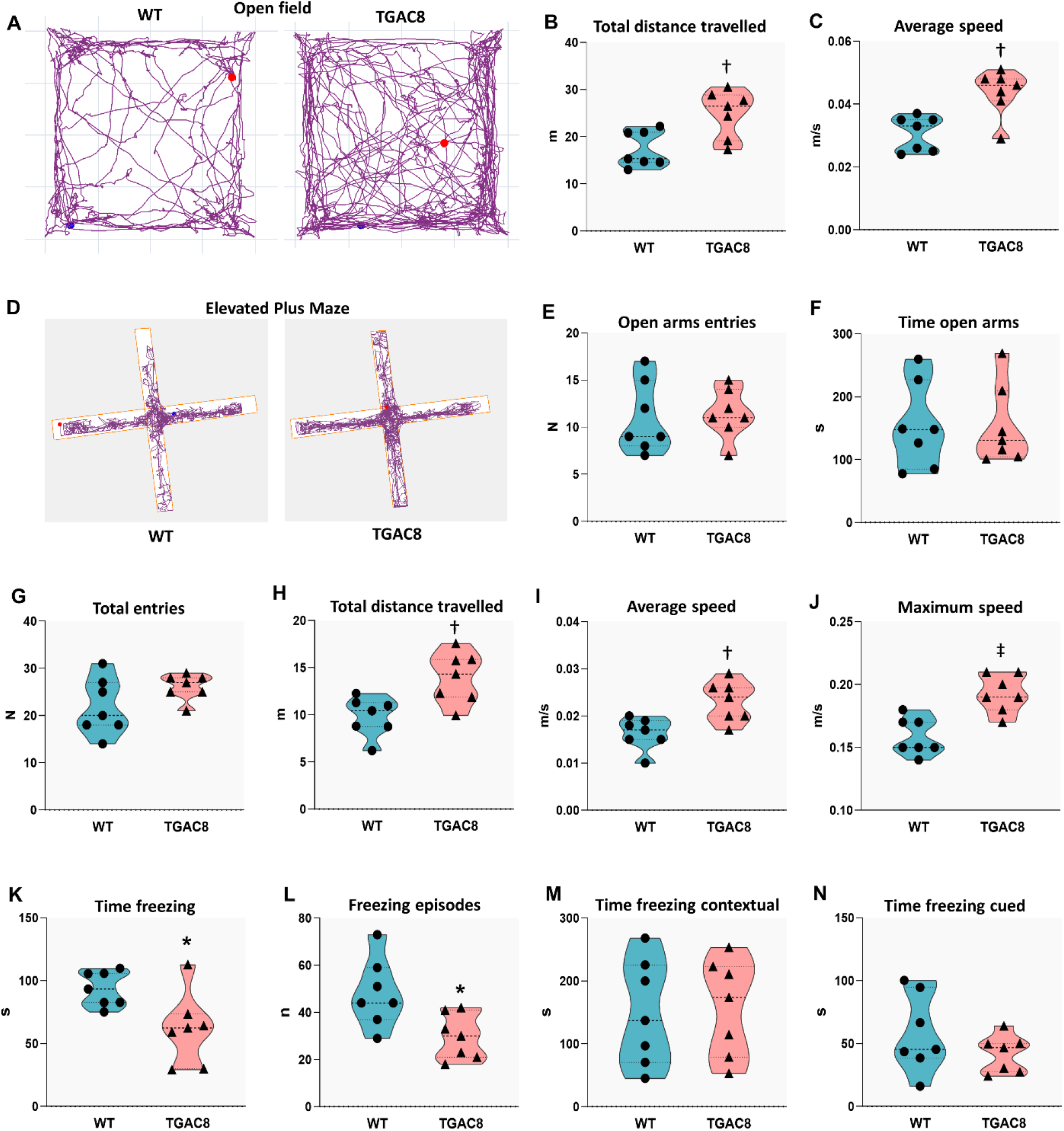
TGAC8 mice display a general increase in locomotor behavior. (A) Plot of the animals movements during OF test, WT vs TGAC8 mouse; (B) Open Field test, total distance travelled; (C) Open Field test, average speed; (D) Plot of the animals movements during Elevated Plus Maze test, WT vs TGAC8 mouse; (E) Elevated Plus Maze test, number of entries in the open arms of the labyrinth; (F) Elevated Plus Maze test, time spent in the open arms of the labyrinth; (G) Elevated Plus Maze test, total entries in the open arms of the labyrinth; (H) Elevated Plus Maze test, total distance travelled; (I) Elevated Plus Maze test, average speed; (J) Elevated Plus Maze test, maximum speed; (K) Elevated Plus Maze test, total time freezing; (L) Elevated Plus Maze test, freezing episodes; (M) Fear Conditioning Test, time freezing in a contextual frame; (N) Fear Conditioning Test, time freezing in a cued frame. Results are shown as violins plot, N=7, unpaired t-test has been performed between WT and TGAC8 groups, *p<0.05, †p<0.01, ‡p<0.001.

### Cardiac AC8 transgenesis increases EEG-gamma frequencies, shifting the movement control from theta to fast rhythms

To gain initial mechanistic clues explaining TGAC8 hyperlocomotion, we implanted dual lead telemetric devices into TGAC8 and aged-matched WT littermates. This approach enabled us to monitor EEG and ECG activities simultaneously (time series), as well as actigraphy (Fig. 2A). The 24 hours recording of the activity confirmed that transgenic mice spent more time moving (active) compared to control littermates (Fig. 2B). Then, we focused on the ECG traces recorded during the mice’s active state. Consistent with previous findings (5, 6), overexpressing AC8 only in cardiac cells increased heart rate and contractility while rendering the heart unresponsive to autonomic nerve system (ANS) surveillance (7). Here, however, we validated and expanded this evidence. Indeed, we observed that TGAC8 mice persistently elevated chronotropic activity paired with a marked drop in both time domains and high- and low-frequency domains of HR variability. The latter was recorded by an ECG branch of the double telemetry implant (p<0.001*** TGAC8 vs. WT) (Fig. 2D-F). Additionally, when examining the active portion (animals moving) of the 24-hrs EEG traces, we discovered that the TGAC8 mice had an overall increase in gamma activity, i.e., *y*1/2/3, spanning from 30 *Hz* to 160 *Hz* (Fig. 2G).

**Figure 2.**
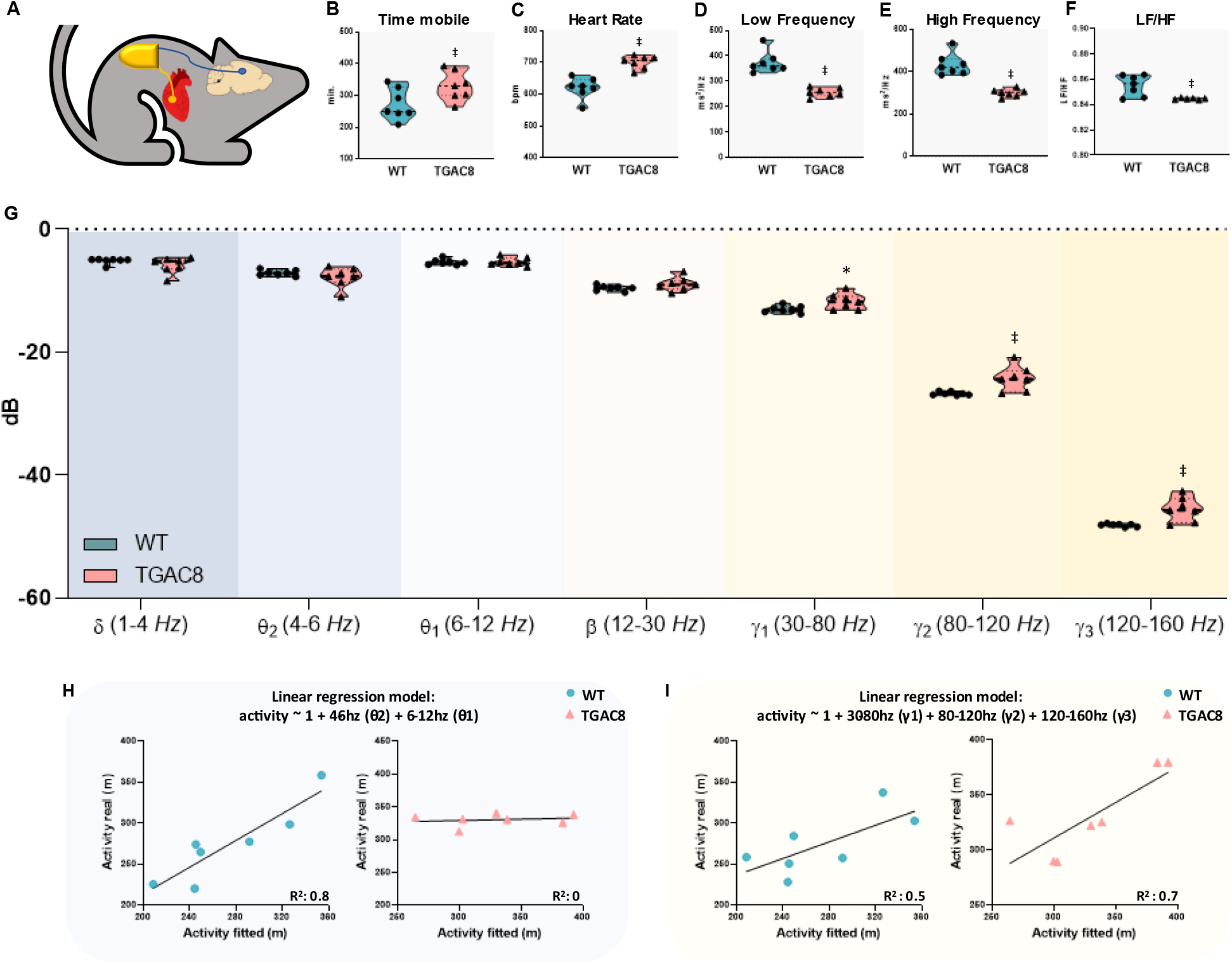
Double ECG and EEG recording, Heart Rate Variability is impaired, while EEG-gamma rhythms are upregulated in TGAC8 mice. (A) Schematic of F20-EET double implant for telemetry recording in mice; (B) Actigraphy, time mobile; (C-F) Heart Rate Variability parameters measured in vivo from telemetric ECG recording: (C) Heart Rate; (D) Low Frequency; (E) High Frequency; (F) Low Frequency/High Frequency ratio; (G) EEG frequencies bands (from δ to γ3) measured in vivo from telemetric EEG recording. Results are shown as violin plots, N=7, unpaired t-test has been performed between WT and TGAC8 groups, *p<0.05, †p<0.01, ‡p<0.001. (H) Linear regression model between activity (time mobile recorded by actigraphy) and θ power; (I) Linear regression model between activity (time mobile recorded by actigraphy) and γ power.

Theta and gamma activities are crucial determinants of overall locomotor behavior in species such as mice (12, 13). Therefore, we next correlated the active movement recorded by the actigraphy with these bands using a linear regression model to determine the degree of association between activity and EEG patterns. We found that the theta bands power (theta-2 and theta-1) is nicely associated with the activity pattern of WT mice (R^2^=0.8), as shown previously (12). Surprisingly, however, the theta band did not influence locomotion in TGAC8 mice (R^2^ = 0) (Fig. 2H). Conversely, the three gamma bands power were reliably associated to active state (moving animal) in TGAC8 mice (R^2^ = 0.7) while maintaining a moderate fit for movements in WT mice (R^2^= 0.5) (Fig. 2I).

In aggregate, these data suggest that cardiac-selective overexpression of AC8 can drive an increase in myocardial performance and concurrently bring about changes in the EEG central patterns, especially within the gamma frequency range. In turn, these modifications can alter the mouse locomotor behavior. Indeed, the current data uncover a shift in the control of locomotion of TGAC8 from the classical theta band to a more prominent role of gamma rhythms.

### The flow of information between the heart and brain is augmented in TGAC8 mice

Parallel to the descending control exerted by the ANS on cardiac function, the peripheral information generated within the heart is communicated to the brain via the intrinsic cardiac nervous system through ascending autonomic fibers (3). We next explored the capacity of the TGAC8 hearts to respond to autonomic input. More in detail, we assessed the intrinsic response of the sinoatrial node (SAN) to external sympathetic and parasympathetic stimuli in isolated atria driven by spontaneous SAN impulses (Fig. 3A). Despite a substantial decreased isoproterenol response (Fig. 3B), the parasympathetic effects evoked by carbachol administration were intact (Fig. 3C). Next, we evaluated the baroreflex *in vivo* via phenylephrine administration. Again, both WTs and TGAC8s responded to the stimulation similarly (Fig. 3D). Thus, we confirmed that, despite some alterations, the autonomic communication between the heart and the brain is still in place in TGAC8 mice.

**Figure 3.**
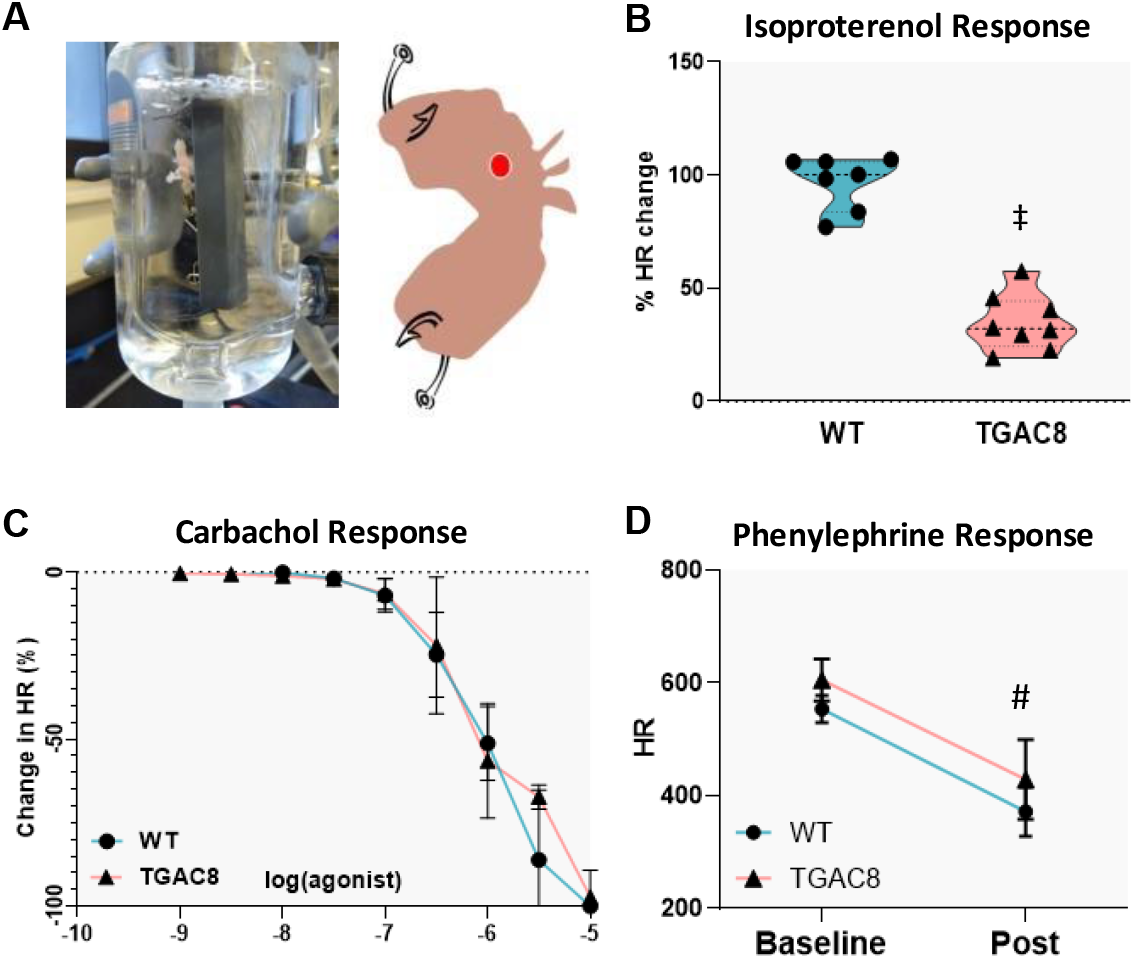
TAC8 mice show a blunted response to sympathomimetic agents, while maintaining parasympathetic reactivity. (A) Representative picture and diagram of isolated atria preparation; (B) Change (in percentage) of heart rate after Isoprot1e8re9nol administration, n=7-8, data are shown as violin plots, unpaired t-test between WT and TGAC8, ‡p<0.001; (C) Change (in percentage) o1f 9h0eart rate after carbachol administration, n=4-5, log dose-response curve. (D) HR change after phenylephrine administration in vivo, resu1lt9s1are shown as mean and SD, n=3-4, two way ANOVA, Baseline Vs Post #p<0.01

We next determined whether the bidirectional communication, in terms of interaction between ECG and EEG signals, is altered in TGAC8. To this end, we used the Granger causality approach to determine whether and to what extent heart-generated electrical signals causally influence electrical cerebral activities (EEG patterns) and vice-versa (14) (Fig.4A, D).

We filtered the ECG traces for Low (LF) and High Frequencies (HF), which substantially represent the autonomic modulation of heartbeat intervals. Analogously, EEG were also filtered to obtain a signal for each band (from δ up to γ3). The GC analysis takes advantage of the variability expressed over time by both brain and heart rhythm. Indeed, it aims to derive the mutual causal influence between heart rate sources (LF and HF) and EEG band activities, and vice versa. Hence, for both ECG and EEG band filtered signals we considered the local amplitude, i.e. envelopes showed in figures 4 A, and D. In terms of brain-to-heart influence, we observed a significant increase in the causal interaction for each EEG band (except for delta) towards LF-filtered ECG RR intervals (Fig. 4B). This is not surprising when considering the necessity of the brain to modulate the elevated HR and contractility of TGAC8 negatively. However, no significant changes were observed in the HF-filtered ECG RR intervals range. Concerning the relation between LF-filtered ECG and EEG, we observed a marked rise in the exchange of information for all the three gamma bands, i.e., the LF component of heartbeat influences and guides changes in TGAC8s EEG patterns in the range of gamma (Fig. 4E). However, we noticed no differences between groups when examining HF-filtered ECG (Fig. 4F).

**Figure 4.**
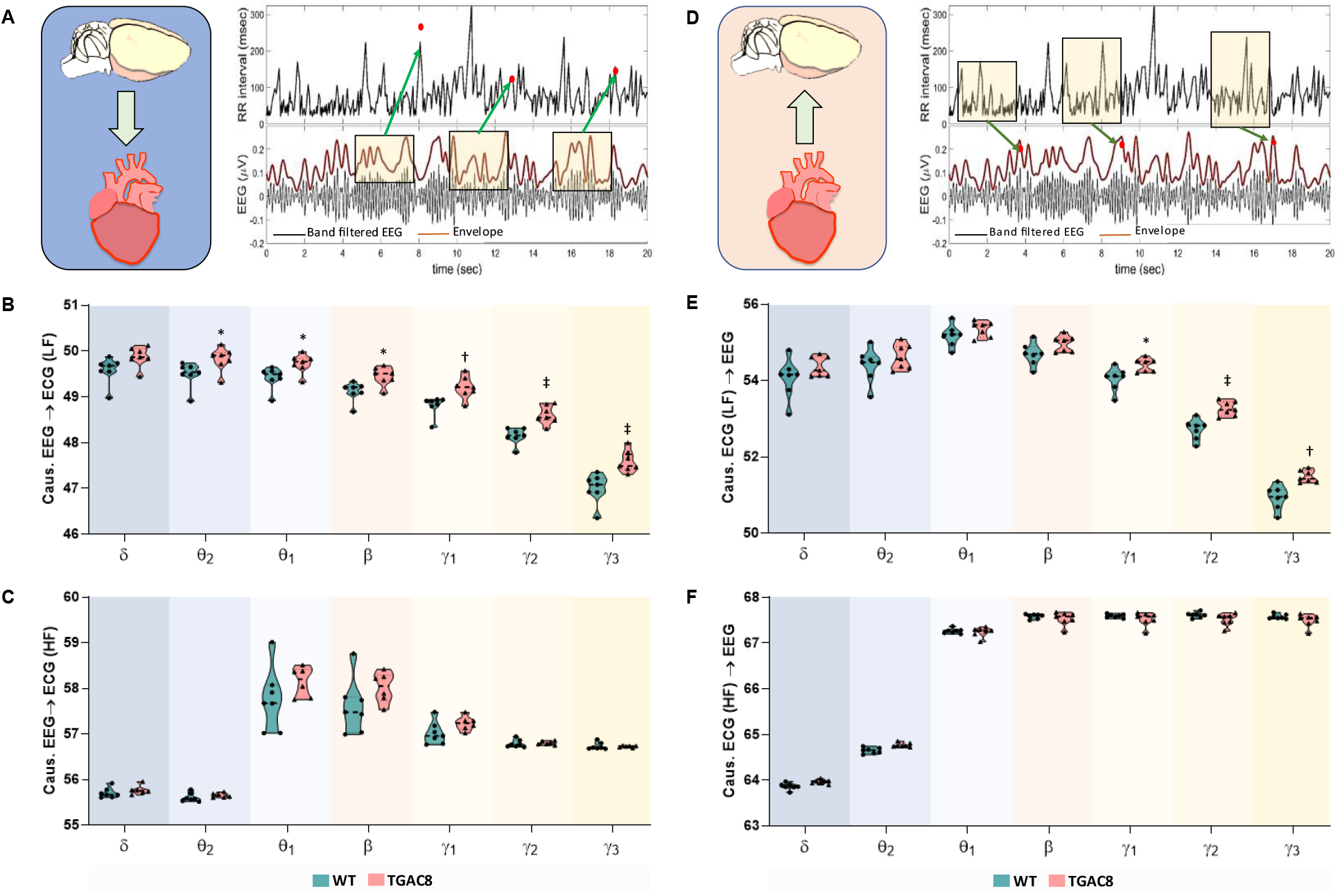
The causal interaction between heart and brain is increased in TGAC8 mice. (A) Granger Causality (GC) brain->heart estimate. The amplitude of the envelope of the band-filtered EEG signal (local amplitude) was extracted and used to estimate its contribution to the prediction of heart rate variability (HRV). The box over the envelope signal depicts the length of past values in the time series used in the GC analysis to predict the current HRV series. To derive the GC brain->heart, this predictive contribution was compared with that made by past values of the HRV series on its own current value; (D) GC heart->brain estimate. HRV time-series (filtered for LF and HF) was extracted and used to estimate its contribution in the prediction of the local amplitude of band-filtered EEG signal. The box over the HRV time series depicts the length of past values in the time series used in the GC analysis to predict current band amplitude. To derive the GC heart->brain, this predictive contribution was compared with that made by past values of the band amplitude series on its own current value; (B-C) dispersion within groups of GC brain->heart estimates both for LF-filtered ECG RR intervals and HF-filtered ECG RR intervals; (E-F) dispersion within groups of GC heart->brain estimated both for LF-filtered ECG RR intervals and HF-filtered ECG RR intervals. Results are shown as violin plots, N=7, unpaired t-test has been performed between WT and TGAC8 groups, *p<0.05, †p<0.01, ‡p<0.001.

We interpret these data to indicate that the flow of information between the heart and brain in TGAC8 is increased. This increment is particularly evident between the LF component of the ECG and the gamma frequencies of the EEG, likely accounting for the gamma rhythms rise observed in TGAC8.

### Changes in GABA and glutamate receptors account for increased gamma activity in the hippocampus of TGAC8 mice

Because EEG recording indicated that TGCA8 mice displayed a marked increase in gamma power -from 30 *Hz* to 160 *Hz* – when these mice were active (Fig.2 G), we next determined what long-term neurochemical adaptations may subtend these changes. The hippocampal formation is one of the primary hubs of gamma activity generation, especially high-speed gamma ripples (15). The earliest model of gamma oscillations generation is based on the reciprocal connections between pools of excitatory pyramidal neurons and inhibitory interneurons, i.e., resulting from a GABAergic/glutamatergic interplay (16). Against this background, we performed a transcriptomic/proteomic analysis to determine eventual changes (long-term adaptations) in the transcription and expression of GABA and glutamate receptors’ transcription and expression levels. The receptor GABA-A is now recognized to play a significant role in gamma activity generation (16). Consistent with this acquired evidence, we found the increased EEG gamma frequency bands in TGAC8 transgenic group coupled with a marked up-regulation of GABA-A. Moreover, the GABA-B, known to inhibit gamma wave generation (17) were downregulated in TGAC8 (Fig.5B). There is a close relationship between GABA-dependent signaling and glutamate receptors synapses in gamma rhythms generation (16, 18). Accordingly, the excitatory receptor mGLU1/5 was upregulated in TGAC8. In contrast, the inhibitory mGLU4/6 and mGLU4/8 were both downregulated in these animals (Fig. 5C). Taken together, these data link the augmented EEG gamma bands to long-term adaptation/remodeling of the GABA/glutamate receptor signaling involved in gamma wave generation, at least at the hippocampal level.

**Figure 5.**
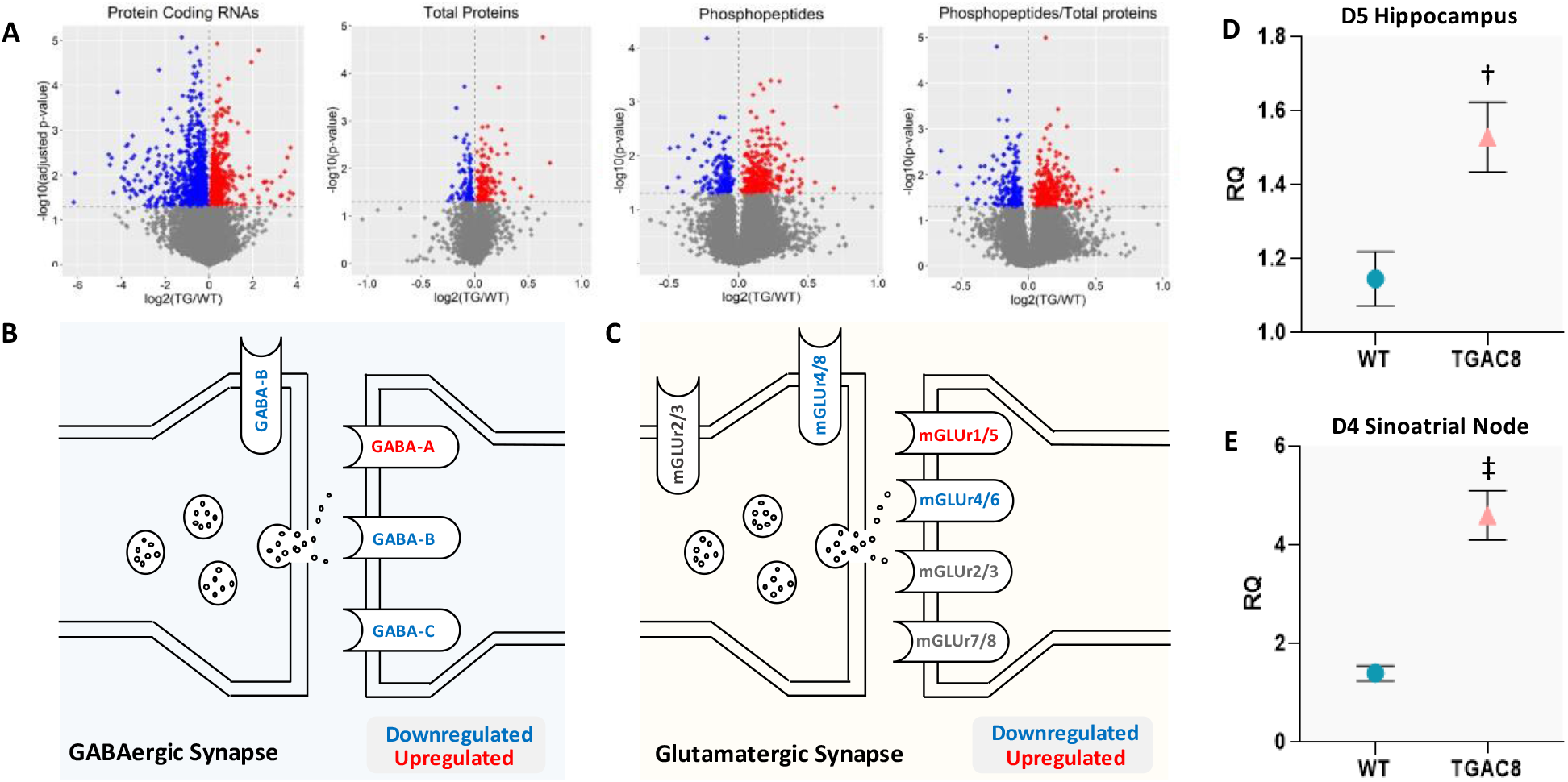
Transcriptomic/proteomic analysis of WT and TGAC8 hippocampus and SA node. (A) vulcano plots of transcriptomic, proteomic and phosphoproteomic analysis of TGAC8 mice hippocampus: protein coding RNAs, proteins, phosphopeptides, proteins/phosphopeptides ratio downregulated in blue, upregulated in red; (B) GABAergic synapse scheme, upregulated receptors in red, downregulated in blue; (C) glutamatergic synapse scheme, upregulated receptors in red, downregulated in blue; (D) Dopamine receptor 5 (D5) transcrption levels in the hippocampus; (E) Dopamine receptor 4 (D4) transcrption levels in the SA node. Results in D and E are shown as mean and SD, n=4, unpaired t-test between WT and TGAC8, †p<0.01, ‡p<0.001.

### TGAC8 mice show an increment of dopamine signaling within the heart, circulating plasma, and hippocampus

The sinoatrial node of the TGAC8s mice displays elevated tyrosine hydroxylase (TH) expression levels with reduced transcription of dopamine beta-hydroxylase: effects converging to augment dopamine bioavailability, as previously shown (7). Moreover, the same study demonstrated a sizable rise in circulating L-Dopa and dopamine in TGAC8 (7). Here, we assessed dopamine receptor transcription levels in the sinoatrial node cells and observed a marked increase in the transcription of the D4 receptor (Fig. 5E). Of note, at the cardiac level, this isoform modulates the activity of the autonomic nervous system (19). This evidence fits very well with the altered bidirectional heart-brain communication detected in the Granger causality analysis. On these grounds, and bearing in mind that DOPA can pass the blood-brain barrier, we hypothesized that increased levels of cardiac-derived DOPA coupled with altered *bottom-up* autonomic signaling account, at least partly, for the hyperlocomotion manifested by the TGAC8 mice. We evaluated dopamine receptor levels in the hippocampus to explore this hypothesis further. Indeed, recent studies proved that the interaction between GABA-A and dopamine D5 receptors is critical to evoking a rise in GABAergic inhibitory postsynaptic currents (IPSCs) (20), an event at the foundation of gamma rhythms generation. In keeping with our initial hypothesis, we found that the expression of D5 receptors was increased in the TGAC8 hippocampus (Fig. 5D).

## Discussion

The central nervous system continuously monitors peripheral internal (and external) body changes (1); for example, the degree of contraction of visceral muscles, heart rate, and metabolite levels in the internal milieu. Collected by the interoceptive system, these modifications signal back to sensory CNS regions encrypting body region representations to command physiological reactions. The latter ultimately modifies body functions to cope with stress conditions and/or maintain organismal homeostasis (1). Our observations fit nicely in this conceptual framework while providing unprecedented experimental evidence that buttresses and expands the somatic theory of emotions (21). Capitalizing on a mouse model that overexpresses AC8 in cardiomyocytes, here we report that TGAC8 mice manifest increased locomotor behavior sustained by enhanced EEG gamma rhythms. In these mice, the analysis of causality interplay (Granger) between ECG and EEG signals attests to a causal involvement of the autonomic component of the heartbeat in shaping EEG gamma oscillations. Mechanistically, the heart of TGAC8 mice showcases heightened dopamine anabolism translating into elevated circulating dopamine/DOPA, paralleled by an upregulation of D4 dopamine receptor. The latter is a primary modulator of cardiac autonomic activity. Centrally, the hippocampus of the TGAC8 mice retains upregulated expression of both GABA-A and D5 dopamine receptors; the activation of the former chiefly accounts for gamma activity generation, while that of the latter exerts a permissive (synergistic) action on GABA-A activation, favoring hippocampal inhibitory currents generations (Fig. 6).

**Figure 6.**
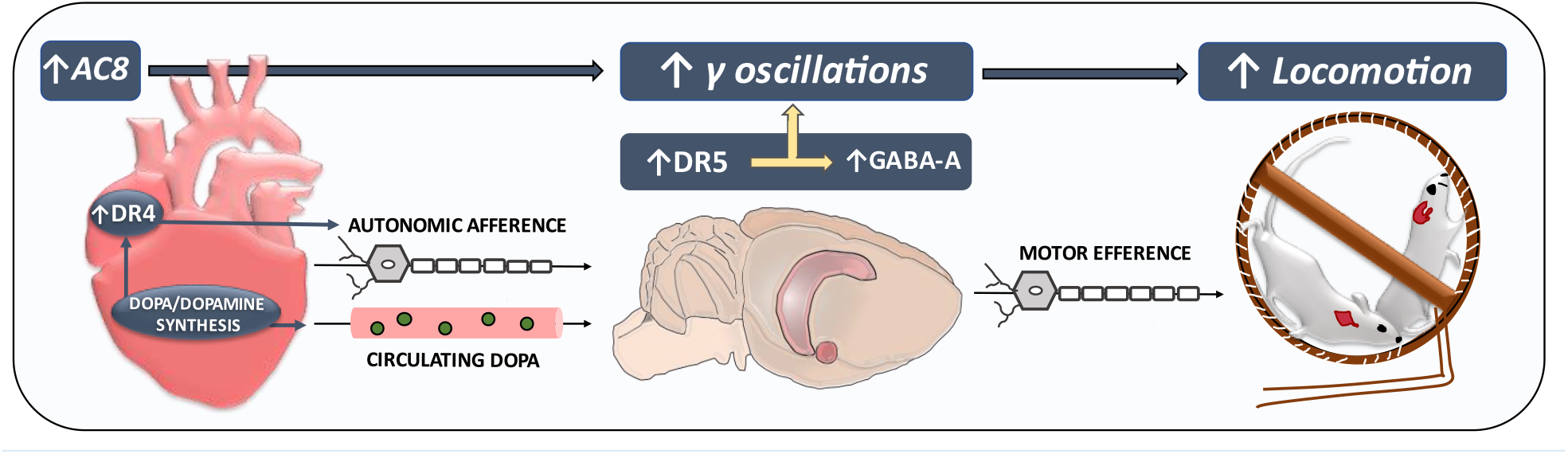
Mechanistic framework of the study. At the cardiac level, mice overexpressing adenylyl cyclase type 8 in cardiomyocytes display heightened dopamine anabolism, thus elevated circulating dopamine/DOPA levels, paralleled by the mRNA transcript upregulation of D4 dopamine receptors - primary modulators of cardiac autonomic activity - in the SA node. Centrally, the hippocampus of the TGAC8 mice retains increased expression of both GABA-A and D5 dopamine receptors. The activation of the former chiefly accounts for gamma activity generation, while the latter exerts a permissive (synergistic) action on GABA-A activation, hence favoring overall hippocampal gamma rhythms generation. Owing to this heightened (bottom-up) heart-brain interaction (confirmed by the Granger causality analysis of the ECG and EEG traces), AC8 transgenesis at the myocyte levels initiates and substains enhanced locomotor activity.

### Overexpressing AC8 in cardiomyocytes triggers a locomotion status akin to perpetual exercise

Mice overexpressing AC8 in cardiomyocytes (TGAC8 mice) exhibit a global and persistent increase in locomotor activity independent of changes in other behavioral variables, such as anxiety. We ground our conclusion on the following experimental evidence: 1) actigraphy (24 hrs. recording) showing increased activity in TGAC8s; 2) open field and elevated plus maze showing increased moving time and distance traveled; 3) superior motor performance, i.e., higher average speed and higher maximum speed and decreased freezing. Of relevance, both OF, EPM, and light-dark box reveal the absence of anxiety-like behavior in these mice. This conclusion is reinforced by the fear-conditioning test, an anxiety-centered test completely independent of motor influences.

We advance the possibility that the rise in locomotor activity found in the TGAC8 is “functionally justified” and congruent with chronic peripheral physiological modifications, such as the persistently elevated heart rate. Put simply, the TGAC8 mouse heart experiences and must handle a status akin to “perpetual exercise” where the brain, in keeping with the somatic theory of emotions, adapt its outflow, switching the animal behavior from a state of rest to an active one, such as running. This scenario also corresponds to the significant cardiac workload sustained by the TGAC8 mice. Moreover, that these transgenes showcase hyperlocomotion but not anxiety is, in our opinion, not so surprising. Indeed, their cardiac chronotropic rhythm is regular and continuous, mimicking a “perpetual exercise” status. Conversely, sudden and irregular bouts of heart rhythm perturbations typically subtend an anxious behavior, such as palpitations and/or frank arrhythmias (22, 23). We posit that the modified pattern of brain-heart communication observed in the TGAC8 mice occurs in anticipation of exercise performance but in the reverse order. Normally, the brain commands the heart to increase its beating rate, ultimately preparing the body to augment physical efforts (24). In stark contrast, in the present experimental setting, the AC8-driven cardiac humoral (endocrine) and functional alterations signal back to the brain to initiate locomotion. In essence, we postulate that, as with other behavioral domains such as emotion control, a heart-brain loop also exists to start and/or fine-tune locomotion.

### Gamma rhythms, locomotion, and heart-brain causal interaction

In rodents, hippocampal theta and gamma rhythms have been linked to locomotion, controlling both its initiation and speed (12, 13). Heightened running speed is accompanied by significant, systematic rises in the frequency of hippocampal network oscillations spanning the entire gamma range (30–120 *Hz*) and beyond (13). Changes in frequency correlate strongly with firing rate changes of individual interneurons mediated by GABA-A receptors, consistent with gamma generation models (16). In our experimental setting, the 24 hr. EEG recordings documented that, during the active phases, the TGAC8 mice experienced a marked rise in gamma frequencies from 30 *Hz* to 160 *Hz*. When correlating the animal motor activity with theta and gamma frequencies, we found a prominent correlation between theta frequencies and movement in WTs. Conversely, such correlation was no longer evident (completely abolished) in TGAC8 mice where gamma activity correlated with locomotion.

Delving further into this finding, we performed some transcriptomic and proteomic analyses of the hippocampus, i.e., the fundamental hub of gamma activity generation and a primary player in locomotor activity control (12, 15). We observed a biomolecular profile that nicely mirrors what was detected at the EEG level. Indeed, all receptors accounting for gamma activity generation (16, 18) were upregulated (especially GABA-A and mGLUr 1/5), while those implicated in gamma inhibition (17), i.e., GABA-B and mGLUr 4/8, were down-regulated.

Finally, to establish a causal link between the EEG pattern changes and cardiac activity stemming from AC8 cardiac overexpression, we evaluated the extent and characteristics of the heart/brain bidirectional communication, applying the Granger Causality approach to EEG and ECG traces. The main finding is that the autonomic component of the heartbeat (especially LF) causally influences the gamma frequencies in a *bottom-up modality*. Specifically, our analysis takes advantage of the variability expressed over time by both the heart rhythm (decomposed in LF and HF) and EEG frequency bands and checks whether amplitude variations of the former are predictive of changes in the latter.

Prior functional magnetic resonance imaging studies using the Granger approach revealed the hippocampus as a pivotal structure in the top-down control of the cardiac output. However, the same study failed to identify causal interactions in the opposite direction, i.e., from the heart to the brain, likely because performed in animals under resting conditions (25). However, recent work by Candia-Rivera and collaborators unveiled the temporal dynamics of brain and cardiac activities in human subjects who underwent emotional elicitation through videos. This research demonstrates, via computational modeling, that emotional stimuli modulate heartbeat activity, stimulating a specific cortical response (4). These observations corroborate both Ledoux’s and Damasio’s theories on emotions. However, in the study by Candia-Rivera and colleagues, the original stimulus, i.e., the trigger of the adaptive brain response, was not peripheral (somatic), in principle. In our study, we specifically overexpressed AC8 peripherally in the heart, resulting in heart rate elevation, heart contractility, and alterations in heart and plasma neurotransmitter levels. While escaping top-down sympathetic control, these alterations would modify the heart-brain communication in a mere bottom-up modality.

### Dopamine and the heart-brain loop

Sinoatrial node cells of the TGAC8s mice harbor markedly elevated levels of tyrosine hydroxylase (TH) with reduced transcription of dopamine beta-hydroxylase, as shown previously (7). And these changes lead to elevation of cardiac and systemic DA/DOPA availability (7). Here, we add to this mosaic two new pieces: first, TGAC8 sinoatrial node cells display a marked rise in D4 transcription levels (Fig. 4E); second, these transgenic mice have an increased hippocampal transcription of D5 receptor (Fig. 4D). In aggregate, the new evidence may have the following functional consequences. First, a recent study shows how atrial D4 can modulate vagal activity (19) (i.e., heart-to-brain and brain-to-heart interaction). This fits very well with what we have observed at the level of bidirectional heart-brain communication by Granger analysis. Second, the incremented levels of circulating DOPA output by the TGAC8 heart can permeate the brain-blood-barrier (BBB), reaching central dopaminergic stations fundamental for the generation and maintenance of locomotor activity such as the basal ganglia (26). Third, the D5 receptor, upregulated in the TGAC8 hippocampus, is one of the primary activators of GABA-A, the main catalyst of gamma activity (16). Accordingly, recent work shows that functional cross-talk exists between GABA-A and D5 in hippocampal neurons, favoring inhibitory current generations (19). Thus, the present results support the novel idea that cardiac dopaminergic signaling shapes heart-to-brain communication through direct endocrine mechanisms (systemic DOPA production) and autonomic ones (D4 modulation on afferent vagal transmission), as well as indirect ones, such as hippocampal upregulation of the D5 receptors.

### Limitations and studies in perspective

The chosen experimental model, i.e., persistent intracardiac AC8 overexpression, can be perceived as a limitation of the present study because it is unlikely that AC8 could be so selectively upregulated in a clinical setting. However, such an artifact is instrumental to the current research’s conceptual and methodological strut: to generate a peripheral physiological variation to assess how cardiac autologous humoral/functional signaling reverberates back to the brain. Future studies could use, for instance, a loss-of-function approach instead, determining, for example, the repercussions of cardiac-specific deletion of AC8 on heart properties and locomotion to further validate the current findings. Admittedly, we focused primarily on the hippocampus because it regulates locomotor activity while simultaneously generating gamma EEG activity (9). However, we believe that the results of our study will stimulate future studies focusing on other brain regions/areas that are crucially involved in the directional heart-brain communication (for example, the nucleus tractus solitarius or the hypothalamus) or movement control (e.g., basal ganglia and septum).

## Conclusions

These studies inform how heart-initiated specific humoral/functional changes (in the present case, elevated dopamine/DOPA signaling and increased myocardial performance due to AC8 overexpression) trigger modifications of just as definite brain areas (hippocampus) to shape specific adaptive responses, such as locomotion. These data reinforce and expand the relevance of somatic theories of emotions.

## Materials and Methods

### Animals

All studies were executed in agreement with the Guide for the Care and Use of Laboratory Animals published by the National Institutes of Health (NIH Publication no. 85-23, revised 1996). The experimental procedures were approved by the Animal Care and Use Committee of the National Institutes of Health (protocol #441-LCS-2016). A breeder pair of TG^AC8^ mice, generated by ligating the murine α-myosin heavy chain promoter to a cDNA coding for human AC8 (27), were a gift from Nicole Defer/Jacques Hanoune, Unite de Recherches, INSERM U-99, Hôpital Henri Mondor, F-94010 Créteil, France. Wild type (WT) littermates, bred from the C57BL/6 background, were used as controls.

### Behavioral tests

All the behavioral tests were conducted on mice with transmitters already implanted. Total time mobile or immobile was evaluated during the 24 hours of EEG/ECG recording phase through actigraphy (included in the double implant). All the other behavioral assays were conducted adopting a one-week interval between tests to allow the mice to recover. Tests were performed at the same time of day, with no reversed dark/light cycle.

### Open field

Open field (OF) is a gold standard behavioral test used to study locomotor activity, exploratory behavior, and anxiety-like behavior in rodents (28). The animal is free to explore a circumscribed environment (50cm x 50cm arena surrounded by walls) for 10 minutes, and its locomotor activity and tendency to explore open spaces or hide in the peripheral corners of the arena is evaluated. The analysis of the behavioral parameters has been performed using ANY-maze software (Stoelting Co., IL, USA).

### Elevated plus maze

Elevated plus maze (EPM) is another consolidated test to study rodents’ anxiety-like behavior and locomotor activity. Elevated Plus Maze test exploits the rodent’s conflict between aversion to open spaces (i.e., hides in closed arms of a labyrinth) and instinct to explore new environments (e.g., exploration of open arms of the same maze). The testing apparatus consisted of four white arms at a 90° angle from each other. Tall walls surrounded two “closed” arms, and two “open” arms had no walls (28). Animals were placed at the center of the apparatus facing an open arm and allowed to explore for 5 minutes freely. The analysis of the behavioral parameters has been performed employing ANY-maze.

### Light-Dark Box

The light/dark box (LDB) test is based on the innate aversion of mice to illuminated spaces and the spontaneous exploratory activity in response to minor stressors, such as a new environment and light. This experimental task allows the evaluation of the level of anxiety-like behavior experienced by the animals. The test apparatus comprises a small, dark, and larger illuminated compartment (29). The analysis of the behavioral parameters has been performed through ANY-maze software.

### Fear Conditioning Test

The fear conditioning (FC) test measures the capacity of rodents to learn and recall an association between environmental cues and aversive experiences. Testing was conducted in standard conditioning chambers (*MED-Associates; St Albans, VT*) equipped with a stainless-steel rod floor that delivered an aversive shock, a ventilation fan emitting background white noise, video recording camera, and a white light that illuminated the chamber. During the 8-minute training session mice were placed in the training context and allowed to explore the environment freely. After two minutes, an auditory cue (80 dB) was presented for 30s, with an electric footshock (0.35 mA) delivered continuously during the final 2s of the auditory cue. The tone-shock pairing was repeated on two separate occasions (at 240s and 360s). Mice were left undisturbed in the chamber for 90s and then returned to the home cage to conclude training. Approximately 24 hours later, the conditioning acquisition mice were individually retrieved and placed in the same training context for five minutes for contextual conditioning testing. One hour later, mice were tested for cued conditioning. In this paradigm, mice were first acclimated to a new room for one hour, with red room lighting. Cued fear conditioning was measured in a different context that utilized a white foam floor covering the electrical grid, a modified chamber shape from a black triangle insertion, and white noise was absent. During testing, mice were exposed to the same auditory tone for 3 minutes after an initial habituation period of 3 minutes. Freezing behavior during the test was recorded as an index of associative learning (11).

### Telemetry double implant to simultaneously monitor the EEG and ECG

Telemetric radio transmitters (F20-EET; Data Sciences International (DSI), St. Paul, MN) were surgically implanted in young (3-4 months) WT and TG^AC8^ mice as described (30). Briefly, two surface electrodes, a positive electrode (parietal cortex; AP, -2.0 mm; L, 2.0 mm) and a reference electrode (cerebellum; AP, -6.0 mm; L, 2.0 mm), were passed subcutaneously to the cranial base and placed directly on the dura mater. Two additional bipotential electrodes were routed subcutaneously via a vertical midline incision overlying the abdomen with leads situated in the right upper chest and lower left abdomen below the heart to monitor continuous heart rate (Fig.1). Following a two-weeks recovery period, 24-hr free behaving electrocardiogram (ECG) and electroencephalogram (EEG) were recorded at a sampling rate of 1000 Hz and 500 Hz, respectively, with simultaneous activity recording using the Dataquest ART acquisition system (DSI, version 4.36). Average activity counts were obtained every 10s.

### Signal analysis

#### ECG analysis

Scoring of wake-sleep states (NeuroScore, DSI, version 3.2.0) allowed for selecting all 10 seconds wake epochs and extracting signals.

After appropriate pre-processing (see Supplemental Information), the series of heartbeat time intervals (RR series) was processed to derive HRV features. All features were signal-length independent (31) estimated on 10 seconds epochs to account for highly frequent wake/sleep switches, thus describing the short-time heartbeat dynamics. More in detail, we considered: the mean value of RR intervals and, from frequency spectrum analysis (Hamming-windowed FFT), the power in low frequency (LF: 0.15–1.5 Hz) and high frequency (HF: 1.5–4 Hz) bands. We also estimated some nonlinear features that provide additional information about cardiac dynamics.

According to previous works studying non-linearity in a heartbeat series (7), we characterize the RR series auto similarity through the Detrended Fluctuation Analysis (DFA).

#### EEG Spectral analysis

For each 10 seconds wake epoch, a spectral analysis was performed on the EEG signal. The mean power spectrum density characterizing the wake state of each animal was evaluated by applying a Hamming-windowed Fast Fourier Transform (FFT) and averaging them. Power values were measured in dB, thus, data from FFT were log-transformed (32).

#### Brain/heart communication

The brain/heart communication study was based on a Granger Causality (GC) analysis. GC_x->y_ is a measure of the contribution of the past of the x(t) time series to the prediction of the present value of y(t), compared to the contribution of the past of the *y(t) time series* in the prediction of its own present value. In our application (Fig. 4) x(t) and y(t) are signals related to brain and heart functioning, and the GC has allowed estimating heart rhythm influence on brain EEG rhythms and vice versa (by switching x and y). Specifically, our analysis takes advantage of the variability expressed over time by both the brain rhythms and heart rhythms and checks whether amplitude variations of the former are predictive of changes in the latter or vice versa. The GC analysis was conducted using the Causal Connectivity Toolbox (33), which assumes a linear model and order within the model, estimated using the Akaike Information Criterion. Herein, in unison with the GC applicability criteria, EEG epochs satisfying the stationarity test (34) were retained for the analysis (more than 95% retained). For each mouse, the median of their epochs model orders was chosen as the representative order. The 95^th^ percentile of ‘animals’ model orders was taken as the overall order (14): from data collected in this experiment, this corresponded to 20 samples, equivalent to 2 seconds. The overall model order was applied to the GC estimates of all wake epochs for all mice (both groups). Finally, the validity of the model order was verified by estimating the model consistency (percentage of data correlation structure explained by the model)^19^. Consistency values higher than 75% were considered satisfactory, and all periods had mean consistencies of 90% or above.

#### RNA sequencing (RNA-seq) and transcriptomic data analysis

We used six hippocampal formations from six transgenic mice and six hippocampal formations from six wild-type littermates for the RNA sequencing experiment. The differential expression gene (DEG) analysis was performed using the DESeq2 package in R language. In total, the expression of 17542 protein-coding mRNAs was identified. The adjusted p-value and fold change calculated by DESeq2 (35) were used to draw a volcano plot to figure out the expression changes in transcriptome. Specifically, 927 genes were downregulated (-log10(adjusted p-value) > 1.3 and log2(fold change) < 0) and 666 upregulated ((-log10(adjusted p-value) > 1.3 and log2(fold change) > 0).

#### Mass spectrometry and proteomic and phosphoproteomic data analysis

4 hippocampla tissues from 4 transgenic mice and 4 hippocampal tissues from 4 wild-type littermates were used for the mass spectrometry experiment. Samples were labeled with the 10-plex tandem mass tag (TMT) according to Thermo Scientific’s TMT Mass Tagging kits protocol. About 5% of the labeled tryptic peptides from each of the 24-fractions were used for global proteomics analysis, while the remaining 95% in the 24-fractions were then pooled into 12-fractions and were subjected to subsequent TiO2-enrichment with a Titansphere Phos-kit (GL biosciences Inc.). All MS and MS/MS raw spectra of TMT experiments from each set were processed and searched using Sequest HT algorithm within the Proteome Discoverer 2.2 (PD2.2 software, Thermo Scientific). The subsequent analyses were adapted from our previous work (36). Specifically, the raw counts of total proteins were normalized to the total counts in each sample, and log2 was transformed. At last, p-values were calculated from the normalized and transformed counts, and fold changes between TG and WT were calculated upon the average expression between the two groups. For quantitative phosphopeptides analysis, additional phosphorylation on Ser, Thr, Tyr residues were specified as variable modifications. We performed the subsequent analyses in two strategies, one for the phosphopeptides and another for the ratio between phosphopeptides/total proteins (also called normalized phosphorylation). For the for the ratio between phosphopeptides/total proteins part, the raw count of phosphorylation was first divided by the raw count of its corresponding protein to obtain the raw normalized phosphorylation count. Then, the raw phosphopeptides or raw normalized phosphorylation counts were normalized to the total counts in each sample, and log2 transformed. At last, p-values were calculated from the normalized and transformed counts, and fold changes between TG and WT were calculated upon the average expression between the two groups.

The p-value and fold change in total proteins, phosphopepetides, and phosphopepetides/total proteins were used to draw volcano plots to determine the expression changes in proteome and phosphoproteome. Specifically, 84 proteins were downregulated (-log10(p-value) > 1.3 and log2(fold change) < 0) and 123 upregulated ((-log10(p-value) > 1.3 and log2(fold change) > 0); 158 phosphopeptides were downregulated (-log10(p-value) > 1.3 and log2(fold change) < 0) and 286 upregulated ((-log10(p-value) > 1.3 and log2(fold change) > 0); 196 ratios of phosphopeptides to proteins were downregulated (-log10(p-value) > 1.3 and log2(fold change) < 0) and 290 upregulated ((-log10(p-value) > 1.3 and log2(fold change) > 0). The normalized and transformed counts and fold changes between TG and WT were calculated upon the average expression between the two groups. For quantitative phosphopeptides analysis, additional phosphorylation on Ser, Thr, Tyr residues were specified as variable modifications. We performed the subsequent analyses in two strategies, one for the phosphopeptides and another for the ratio between phosphopeptides/total proteins (also called normalized phosphorylation). For the for the ratio between phosphopeptides/total proteins part, the raw count of phosphorylation was first divided by the raw count of its corresponding protein to obtain the raw normalized phosphorylation count. Then, the raw phosphopeptides or raw normalized phosphorylation counts were normalized to the total counts in each sample, and log2 transformed. At last, p-values were calculated from the normalized and transformed counts, and fold changes between TG and WT were calculated upon the average expression between the two groups. The p-value and fold change in total proteins, phosphopeptides, and phosphopeptides/total proteins were used to draw volcano plots to determine the expression changes in proteome and phosphoproteome. Specifically, 84 proteins were downregulated (-log10(p-value) > 1.3 and log2(fold change) < 0) and 123 upregulated ((-log10(p-value) > 1.3 and log2(fold change) > 0); 158 phosphopeptides were downregulated (-log10(p-value) > 1.3 and log2(fold change) < 0) and 286 upregulated ((-log10(p-value) > 1.3 and log2(fold change) > 0); 196 ratios of phosphopeptides to proteins were downregulated (-log10(p-value) > 1.3 and log2(fold change) < 0) and 290 upregulated ((-log10(p-value) > 1.3 and log2(fold change) > 0).

#### RT-qPCR

RT-qPCR of the hippocampus and SA tissue was performed to determine the transcript abundance of dopamine receptors (n=4 WT and 4 TG^AC8^ mice). RNA was extracted from the hippocampus with TRIzol™ Reagent (Thermo Fisher Scientific, Waltham MA) and DNAse on column digestion. The cDNA was prepared using MMLV reverse transcriptase (Promega, Madison, WI) using 500 ng of total RNA per reaction. RT-qPCR was performed using a QuantStudio 6 Flex Real-Time PCR System (Thermo Fisher Scientific, Waltham MA) with a 384-well platform. The reaction was performed with a FastStart Universal SYBR Green Master Kit with Rox (Roche, Indianapolis, IN) using the manufacturer’s recommended conditions; the sizes of amplicons were verified. Each well contained 0.5 μl of cDNA solution and 10 μl of the reaction mixture. Each sample was quadruplicated. Preliminary reactions were performed to determine the efficiency of amplification. RT-qPCR analysis was performed using the ddCt method. *Hprt* was used as a housekeeping gene and results represented as Mean of Relative Quantification (RQ) normalized to *Hprt* +/- SD (8). Primers were selected with Primer Express 3.0 software (Applied Biosystems).

### Statistical analysis procedures

Results are presented as violin plots. All parametric data were analyzed by unpaired t-tests between the WT and TGAC8 groups. Correlations between groups of values were evaluated through multiple linear regressions (37).

## Supplemental Information

**Figure 1:**
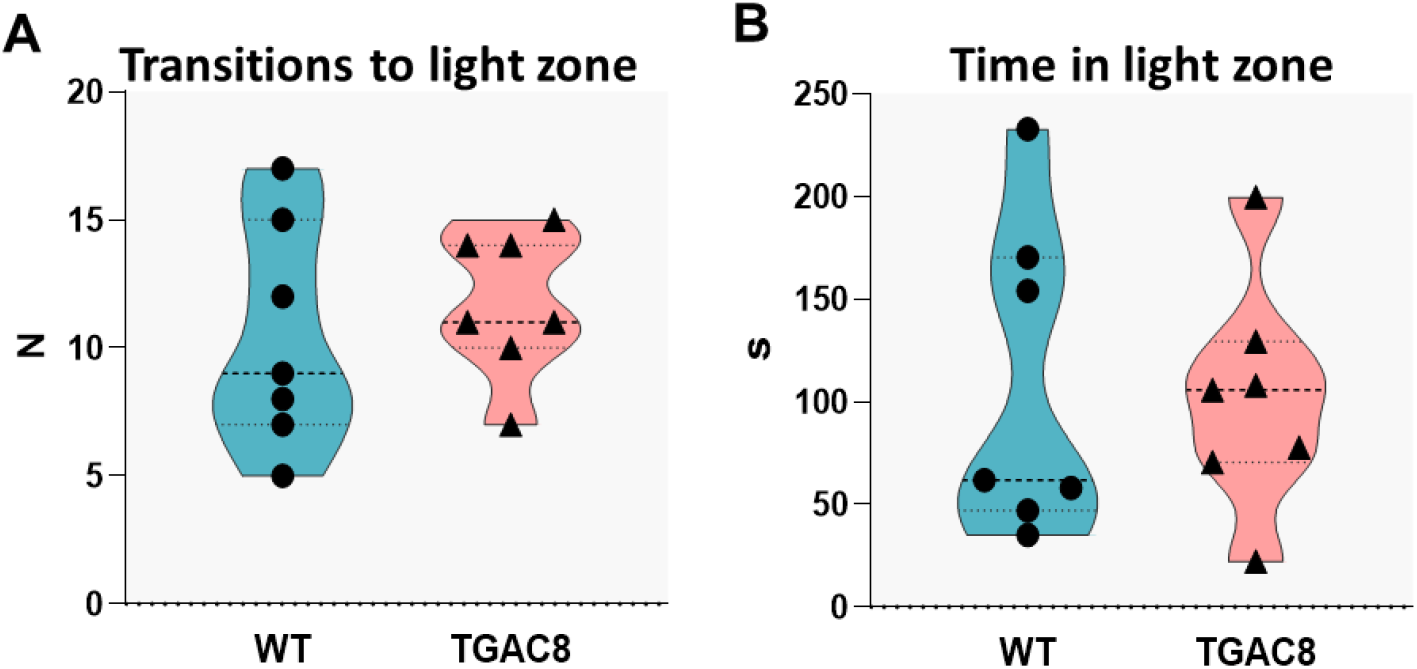
Light-dark box test, TGAC8 mice do not show signs of anxiety-like behavior. A) Light-dark box test, transitions to light; B) Light-dark box test, time in the light zone. Results are shown as violins plot, N=7, unpaired t-test has been performed between WT and TGAC8 groups.

